# Transition between conformational states of the TREK-1 K2P channel promoted by interaction with PIP_2_

**DOI:** 10.1101/2022.02.27.482173

**Authors:** Adisorn Panasawatwong, Tanadet Pipatpolkai, Stephen J. Tucker

## Abstract

Members of the TREK family of two-pore domain (K2P) potassium channels are highly sensitive to regulation by membrane lipids, including phosphatidylinositol-4,5-bisphosphate (PIP_2_). This study used coarse-grained molecular dynamics (CG-MD) and atomistic MD simulations to model the PIP_2_ binding site on both the up and down state conformations of TREK-1. We also calculated the free energy of PIP_2_ binding relative to other anionic phospholipids in both conformational states using potential of mean force (PMF) and free energy perturbation (FEP) calculations. Our results identify state-dependent binding of PIP_2_ to sites involving the proximal C-terminus and we show that PIP_2_ promotes a conformational transition from a down state towards an intermediate that resembles the up state. These results are consistent with functional data for PIP_2_ regulation and together provide evidence for a structural mechanism of TREK-1 channel activation by phosphoinositides.

## Introduction

The TREK subfamily of two-pore domain (K2P) K^+^ channels contribute to the resting membrane potential and electrical activity of a wide variety of cell types and tissues including many within the central and peripheral nervous systems (Natale *et al*., 2021). They are involved in processes such as pain perception, neuroprotection, and anesthesia (Enyedi & Czirják, 2010) and therefore make attractive therapeutic targets (Busserolles *et al*., 2020).

TREK channel activity is regulated by a wide variety of physical and chemical stimuli. This polymodal regulation allows them to integrate cellular electrical activity with a diverse range of cellular signaling pathways. In particular, TREK channels are mechanosensitive and highly sensitive to their lipid membrane environment. Previous studies have shown that membrane tension increases channel activity by expanding the cross-sectional volume of the channel within the bilayer thereby rendering it mechanosensitive (Aryal *et al*., 2017). In addition to the physical properties of the membrane, TREK channels are also highly sensitive to different lipid species as well as both internal and external pH (Maingret *et al*., 1999; Riel *et al*., 2021). For example, amino acids in the C-terminal domain (CTD) are involved in PIP_2_ regulation and intracellular pH sensing (Chemin *et al*., 2005*b*), whilst the CTD also provides binding sites for proteins that can regulate lipid composition such as phospholipase D2 (Comoglio *et al*., 2014; Petersen *et al*., 2019). However, despite the obvious importance of its lipid sensitivity and PIP_2_ regulation in particular, relatively little is known about how PIP_2_ interacts with TREK channels.

Crystal structures of TREK channels have revealed two distinct conformations: an ‘up’ state (PDB entry: 6CQ6) and a ‘down’ state (PDB entry: 4XDJ) (Dong *et al*., 2015; Lolicato *et al*., 2017). The up state has a broader cross-section than the down state within the lower leaflet due to an upwards movement of the M2 and M4 transmembrane helices. However, the structure of the full CTD remains unknown as this domain is truncated in all of the constructs used for crystallization. Both conformations of the channel appear to be conductive (McClenaghan *et al*., 2016; Proks *et al*., 2021) but several studies indicate that the channel becomes more active when it is in the up state (Brennecke & de Groot, 2018; Proks *et al*., 2021).

Movement of the TM helices therefore appears to regulate channel activity, but unlike many other K^+^ channels, this movement does not appear to constrict the pore to provide a simple switch between an open and closed state because TREK channels are gated primarily within their selectivity filter (Bagriantsev *et al*., 2011; Piechotta *et al*., 2011; Lolicato *et al*., 2017). The mechanism by which membrane tension shifts the channel into the more active up state has been studied extensively (Aryal *et al*., 2017; Clausen *et al*., 2017; Rietmeijer *et al*., 2021). However, the mechanisms by which regulatory lipids such as PIP_2_ interact with TREK channels to influence these conformations remains unclear.

In this study we have used both coarse-grained and atomistic molecular dynamics (MD) simulations to model PIP_2_ binding to different conformations of TREK-1 and calculate its energetics of binding relative to other anionic phospholipids. Our results indicate that PIP_2_ promotes a conformational transition towards the up state and proposes a mechanism of activation by PIP_2_ that is consistent with a wide range of evidence from previous binding studies, functional assays and mutagenesis data.

## Results

### Identification of a PIP_2_ binding site

We conducted 10 μs coarse-grained molecular dynamics (CG-MD) simulations to investigate putative lipid binding sites to the up state (PDB entry: 6CQ6) (Lolicato *et al*., 2017) and the down state conformations of TREK-1 (PDB entry: 4XDJ) (Dong *et al*., 2015). The distal C-terminus of the down state structure was extended to match the longer α-helical structure that was resolved in the up state structure (6CQ6). In our simulations, PIP_2_ was initially placed randomly in the lower leaflet with the mixture of PC:PS:PIP_2_ at the ratio of 15:3:2 and simulated for 10 μs for each structure (**Fig 1A**). Similar to previous protein-PIP_2_ interaction studies (Stansfeld *et al*., 2009; Pipatpolkai *et al*., 2020; Duncan *et al*., 2020), we used two criteria to identify possible PIP_2_ binding sites. First, the “residence time” of the PIP_2_ in its binding site was assessed, i.e. the period in which PIP_2_ remained at least 0.6 nm to the residues of interest. Second, we assessed the “occupancy”, defined as the fraction of time that PIP_2_ spends at least 0.6 nm in proximity to a particular residue. These metrics allowed us to cluster any residues with similar residence time and occupancy, therefore defining them as a potential interaction or binding sites.

**Figure 1.**
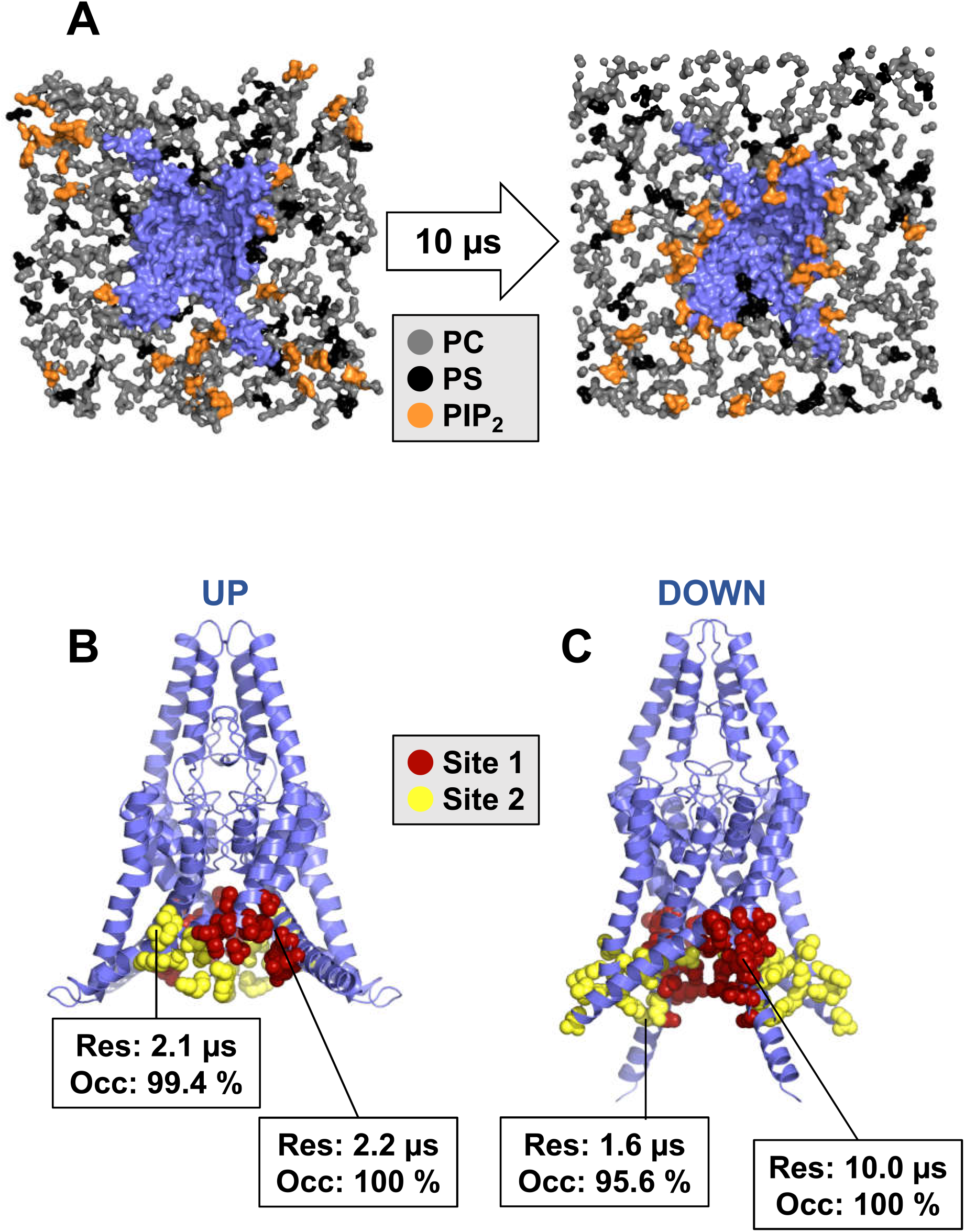
Coarse-grained molecular dynamics simulation. **A**) Schematic representation of the coarse-grained simulation conducted in this study. Only the headgroup of the lower leaflet lipids (PC – grey, PS – black and PIP_2_ - orange) are shown. TREK-1 (blue) in the up state is embedded in the phospholipid bilayer. After a 10 μs simulation (right), PIP_2_ headgroups are distributed more closely at their relative binding site (right) compared to their initial distribution (left). The PIP_2_ binding site on the TREK-1 (**B**) up state or (**C**) down state. The binding site 1 (red) and 2 (yellow) are shown superimposing relative to their crystal structure. Each site is annotated with their residence time within the binding site (Res) and their occupancy (Occ)

We identified two binding sites with occupancy >90% and residence times >2 μs (**Fig 1B,C and Fig S1**). Other sites with residence times less than 0.5 μs were treated as non-specific interactions. In the up state, the two binding sites are adjacent to each other and located between M1 and M4 (**Fig 1B**). In contrast, the binding sites on the down state structure were further away from each other (**Fig 1C**) and the binding site with the highest residence time was similar to the site identified in the up state. However, the second binding site in this conformation is located at the M2/M3 interface, and the two binding sites are separated from each other by the M2 helix. In contrast, these two sites are much closer together when the channel is in the up state (**Fig 1B**).

### PIP_2_ binding affinities between up state and down state

Similar to a previous approach (Corey *et al*., 2019), we next assessed the binding affinity of PIP_2_ in the most prominent binding site using potential of mean force (PMF) calculations to calculate the free energy of binding. We compared the energetic difference between the “bound” state (energy minima) and the “bulk”, where lipids are at least 1.2 nm away from the binding site (**Fig 2A**). We showed that the binding free energy in the most occupied binding sites are −41 ± 3 kJ mol^−1^ and −26 ± 2 kJ mol^−1^ for the up and down states, respectively (**Fig 2B**). Thus, the binding of PIP_2_ appears more favorable in the up state compared to the down state. We also observed a shallower energy landscape in the down state, implying less specificity in the binding site.

**Figure 2.**
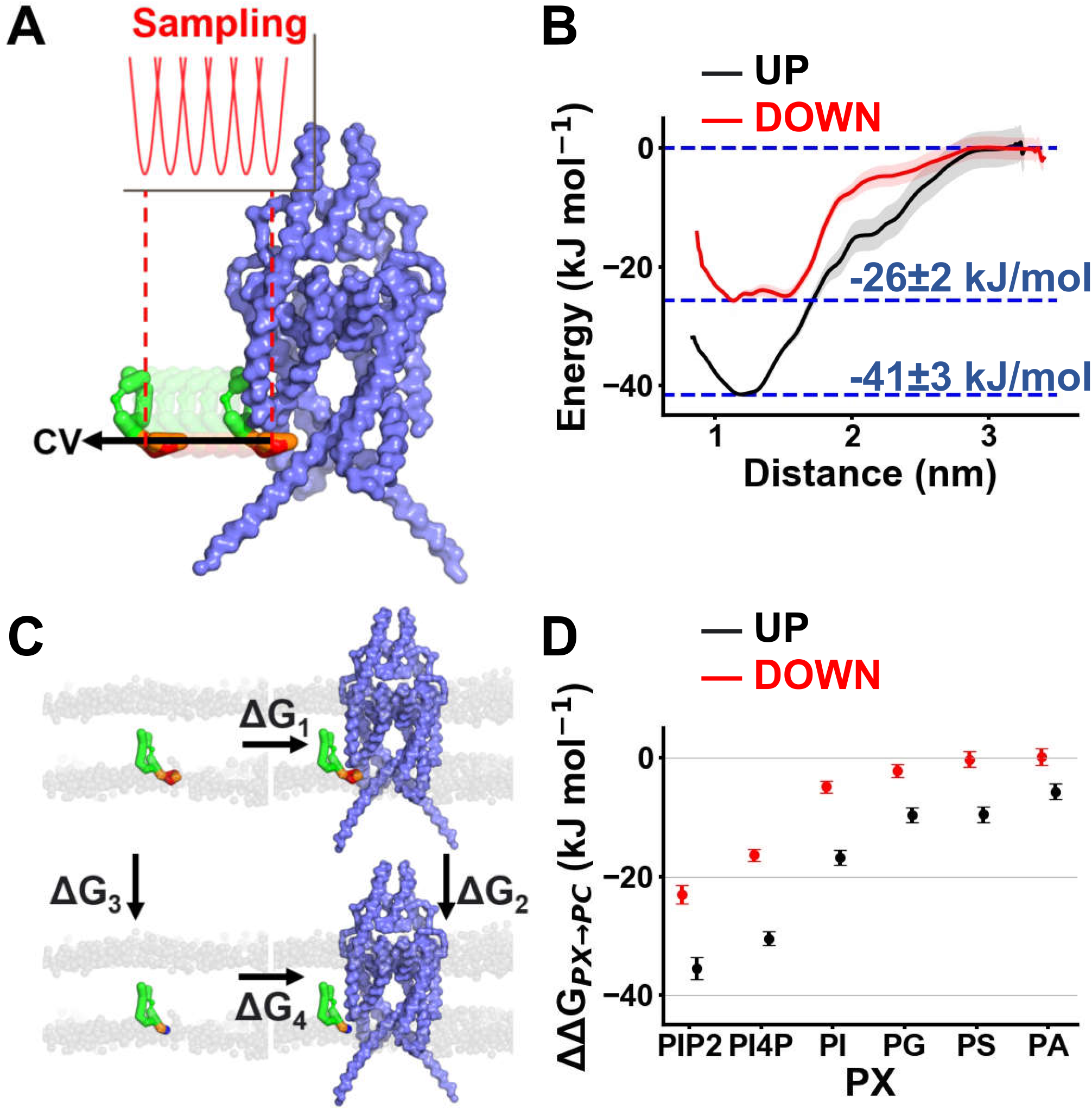
Calculation of PIP_2_ binding free energy relative to other lipids. **A**) Schematic description of the CG-PMF calculations. In this scheme, PIP_2_ (green) in the binding site was steered away from the protein (blue surface) along the collective variable (arrow). The PMF is calculated using umbrella sampling with the force constraint at 1000 kJ/nm^2^/mol (red). **B**) 1D energy landscape calculated from PMF calculation as PIP_2_ is steered away from the binding site from up (black) or down (red) states to the bulk lipid bilayer (PC). Energy landscape is marked at 0 kJ/mol when the PIP_2_ is > 1.2 nm away from the binding site. **C**) Schematic description of CG free energy perturbation (CG-FEP). The PIP_2_ headgroup was alchemically transformed to different lipids (PI4P, PC, PS, PG, PA) in both bound state (ΔG_2_) and in the bulk state (ΔG_3_). This transformation allows relative affinities between two lipids species to be calculated. **D**) Binding free energy of PIP_2_, PI4P, PI, PG, PS and PA relative to PC within TREK-1 PIP_2_ binding site in the up (black) or down state (red). Error bars showed 95% confidence interval calculated based on t-statistics (n = 15).

### Comparing lipid specificity in the primary binding site

To determine the relative binding free energy between PIP_2_ and other lipids in this binding site, we applied the free-energy perturbation (FEP) method to perturb the PIP_2_ headgroup to other lipid headgroups. Relative binding free energies were then calculated relative to PC (**Fig 2C, 2D**). We define the binding free energy of PC to TREK-1 channel to be ~0 kJ/mol as this is the most common lipid in the bilayer. In the up state, the binding free energy difference between PIP_2_ and PC is −36 ± 2 kJ mol^−1^ which agrees well with our PMF calculations (−41 ± 3 kJ mol^−1^). This value is also remarkably close to the value calculated in a fluorescent binding assay which yielded −36 ± 1 kJ mol^−1^ from a dissociation constant (K_d_) of 0.80 ± 0.34 μM (Cabanos *et al*., 2017).

By comparing these binding free energies relative to PC, we observed that channel affinity for PIP_2_ is slightly higher than for PI4P. This suggests the first phosphate group contributes only partially to the binding of PIP_2_. However, the transformation from PIP_2_ to PI showed that the inositol group and the second phosphate group have a much more significant contribution than the first phosphate group. This pattern has also been observed in inwardly rectifying potassium (Kir) channels (Pipatpolkai *et al*., 2020). In addition, we compared the relative binding free energy of PIP_2_ to other anionic phospholipids such as phosphatidylglycerol (PG), phosphatidylserine (PS) and phosphatidic acid (PA) (**Fig 2D**). This showed that the binding free energy of PG, PS and PA are relatively similar (−10 kJ/mol relative to PC).

We next conducted a similar CG-FEP calculation using the structure of the down state TREK-1. Here, we observed a similar difference between the affinity of PIP_2_, PI4P and PI compared to the up state. However, we also observed a much smaller binding energy of PG, PS and PA (~0 kJ/mol relative to PC) in the down state suggesting that any channel activation these anionic phospholipids is only likely to occur in the up state.

Previous study has suggested that PA activates the channel because it competes PIP_2_ away from the binding site. However, our free energy calculation showed that the PA affinity is much lower than PIP_2_ in the up state. Thus, we examined whether PA may accommodate an alternative binding site to the channel, or possibly accumulate near the PIP_2_ binding site. To do so, we prepared a bilayer containing 10% PA and 90% PC and conducted a 10 μs unbiased simulation to identify PA binding to the channel. In the up state, the site where PA binds with the highest occupancy and the longest residence time is not the same site as the PIP_2_ binding site. This PA binding site is located between M1 and M2 helices whereas the PIP_2_ site is located between M1 and M4 helices. At the PIP_2_ site, the occupancy and residence time of PA are much lower than for PIP_2_ (0.46 μs in PA binding site and 2.21 μs in PIP_2_ binding site) (**Fig S2**). This suggests that PA is unlikely to directly compete with PIP_2_ at the PIP_2_ binding site we have identified.

### Atomistic simulation of TREK-1

Full atomistic simulations allow us to observe conformational changes in the M4 helix and can capture the geometry of the binding pocket at a much higher resolution. We therefore took snapshots from the final frame of the CG-MD simulations and used these to seed atomistic simulations. To assess the geometry of the binding pocket, we ran 500 ns simulations of both up and down states with a single PIP_2_ molecule in the binding site, in a bulk PC bilayer (n=3). As TREK-1 is a homodimer, the data collected from each subunit was treated as a single data point (n=6). We defined contact residues as those which spent more than 50% of the simulation time at < 4 Å proximity of the PIP_2_ headgroup.

In the up state, PIP_2_ coordinates with 3 residues on M4 (R297, K301 and K304). These 3 residues are all on the same helical interface of M4 (**Fig 3A**). Within this binding site, R297 and K304 are coordinated between two phosphate groups (P1 and P5) on the inositol ring (**Fig 3A, Fig S3**), whereas K301 is only coordinated to P5. However, the P4 phosphate group is not coordinated by any amino acid residues. These variations may explain the lower relative binding free energy differences when the first phosphate group is perturbed in our CG-FEP calculation.

**Figure 3.**
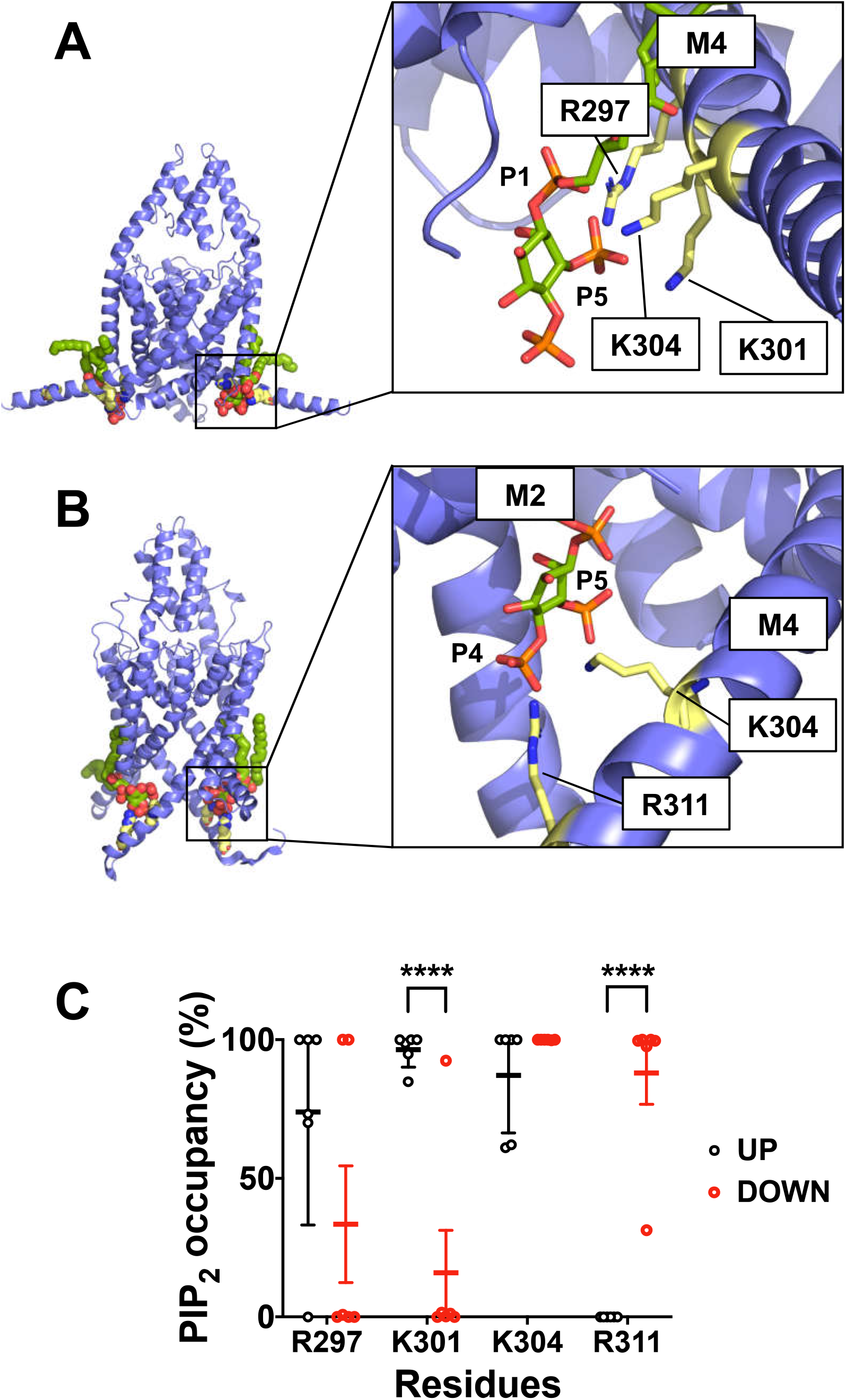
Atomistic simulation of PIP_2_ and TREK-1. Representative interactions between (**A**) up state and (**B**) down state of TREK-1. The backbone of the protein is represented in blue. Key contacting residues contacting to the phosphate group on the PIP_2_ molecule are shown in yellow. **C**) PIP_2_ contact analysis showing the fraction of time that residues are in 4 Å proximity to the PIP_2_ molecule (contact probability) of the up state (black) and down state (red) of TREK-1. Only residues with an average >50% contact probability are shown. Data are collected from three repeats of 500 ns simulations. Error bars showed 95% confidence interval calculated based on t-statistics (n= 6).

In the down state, PIP_2_ only coordinates with 2 residues (K304 and R311). Both are on M4 and form a site that is further away from the pore domain than the binding site in the up state (**Fig 3B**). The phosphate group connected to C4 is also singly coordinated by R311, whereas C5 is coordinated by K304. Interestingly, the C1 phosphate is not coordinated by any amino acid residues within the C-terminus. The fewer contacts made in the down state may therefore explain the lower binding free energy obtained from the PMF calculation.

Comparison of the coordination pattern between the up and down states revealed that PIP_2_ was in contact with K301 more often in the up state compared to the down state (n = 6, Student’s t-test: *P*<0.0001). On the other hand, co-ordination with R311 is significantly more favorable in the down state (n= 6, Student’s t-test: *P*<0.0001) (**Fig 3C**). We then calculated an average number of hydrogen-bond formed in the last 100 ns of the simulation (**Fig S3**). Assuming that the interaction is driven mostly by electrostatic interaction, this suggests that the up state exhibits a greater degree of hydrogen bonding with the key PIP_2_-interacting residues than the down state. This therefore allows us to postulate that PIP_2_ binding is stabilized in the up state, but not in the down state.

### Effect of PIP_2_ on dynamics of the C-terminus

We next investigated the conformational dynamics of TREK-1 in three 500 ns atomistic simulations of both PIP_2_ bound and unbound (apo) states. To assess the protein backbone dynamics, we calculated the root mean square fluctuation (RMSF) on the Cα atom on the protein and compared the effect of PIP_2_ relative to the apo state (control). This difference (ΔRMSF) is shown in **Fig 4A**. This shows that M4 and the C-terminus fluctuates more when PIP_2_ is absent in the down state (ΔRMSF ~ 8 Å) and suggests that PIP_2_ may stabilize the conformation of the C-terminus in the down state. However, the dynamics of the C-terminus are relatively unaffected by PIP_2_ in the up state as the PC headgroups may stabilize the α-helical content of the M4 helix (**Fig 4A**).

**Figure 4.**
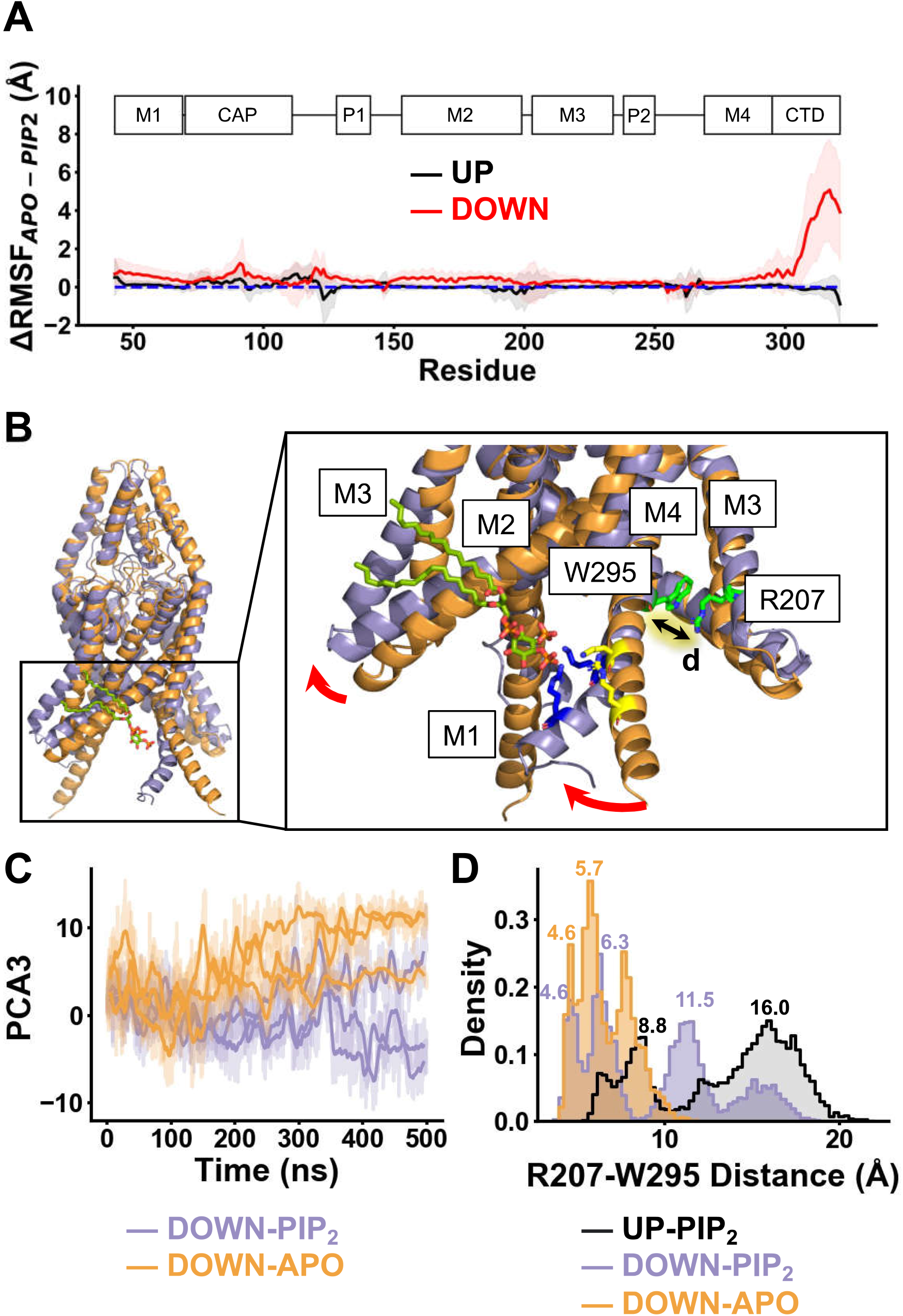
PIP_2_ induces conformational change on TREK-1 C-terminus. A) The difference between the C_α_ RMSF in the presence and absence of PIP_2_ (ΔRMSF) of the up state (black) and down state (red) of TREK-1. The positive ΔRMSF value implies that the backbone is stabilised by PIP_2_. Shaded region represents the propagation of 95% confidence interval. **B**) The conformational change induced by PIP_2_ in the down state. The initial conformation at 0 ns (orange) is compared to the post-simulation at 500 ns (blue) with PIP_2_ (dark green) in the binding site. The key conformational change in the CTD is described with the red arrow. We defined the zipper measurement based on the distance (d) between R207 and W295. Residues co-ordinated with PIP_2_ (K304 and R311) are highlighted before (yellow) and after (blue) 500 ns simulation. **C**) The magnitude of the conformational change along the vector describing the third principal component from the down state plotted against time in the apo state (orange) and PIP_2_-bound state (purple). The darker lines show the running average for each simulation (500 ns) (n = 3). **D**) Distribution plot of the zipper distance collected from the final 100 ns of each simulation for the PIP_2_ bound up state (black), the PIP_2_ bound down state (purple), and apo down state (orange).

### PIP_2_-induced conformational transition

We then focused on the down state where PIP_2_ had large effects on the dynamics of the C-terminus. Over 500 ns, our contact analysis showed that K304 and R311 on M4 bend to interact more closely with PIP_2_ (**Fig 3B, 3C**). This bending enables M2/M3 to move upward. Together, this widens the cavity between M2 and M3 (**Fig 4B**). To quantify the functional dynamics of TREK-1 when PIP_2_ is bound, we calculated the first and the second eigenvector on the protein backbone motion using principal component analysis (PCA). This decomposed the backbone movement relative to the initial structure into orthonormal bases (components). We then projected the movements from the simulations on these components to observe the principal movement ordered by their eigenvalues. The first and second principal components capture the motion of the extracellular cap domain (**Fig S4**) and we focused on the third principal component (PC3), where the conformational change at the C-terminus occurs (**Fig 4C**).

To quantify this change in PC3 (7%), we used the ‘Zipper’ measurement between W295 in M4 and R207 in M3 on the same chain (Aryal *et al*., 2017), and plotted the distribution of this distance (**Fig 4D, Fig S5**). When compared to the up state, where no conformational change occurred at the C-terminus in response to PIP_2_, the transition from the down to the up state involved a re-orientation of W295 in M4. This transition breaks its interaction with R207, which then allows an expansion between TM2 and TM4. We therefore used the distance between the center of mass of R207 and W295 as a metric of the down to up transition (**Fig 4D, Fig S5**) and observed a bimodal distribution, where the majority of the population is distributed between 10 to 20 Å, and a minority at ~ 8 Å.

In contrast, the distance between R207 and W295 is only distributed around 3-9 Å in the down state in the absence of PIP_2_. The down state also displayed a unique population at about 11.5 Å, similar to the up state (**Fig 4D**). This suggests this simulation may capture an initial path of a down to up conformational transition that is driven by PIP_2_.

## Discussion

This study used CG-MD simulation to predict PIP_2_ binding sites on TREK-1 and highlighted differences in PIP_2_ binding between up and down states of the channel. We also quantified the binding free energy of different lipids in both conformations and used principal component analysis to examine the conformational changes induced by PIP_2_. Our results not only identify key residues involved in PIP_2_ binding but also indicate that PIP_2_ induces a conformational transition from the down to the up state.

Although PIP_2_ has a clear activatory effect on TREK channels (Chemin *et al*., 2005*a*, 2005*b*; Lopes *et al*., 2005; Soussia *et al*., 2018; Rivas-Ramírez *et al*., 2020; Riel *et al*., 2021), inhibitory effects of PIP_2_ have also been reported (Chemin *et al*., 2007; Woo *et al*., 2016, 2018). In addition to PIP_2_, other anionic phospholipids such as PA have also been shown to increase channel activity (Chemin *et al*., 2005*b*) and in one particular study (Woo *et al*., 2016) PIP_2_ was shown to have a concentration-dependent biphasic effect on the channel, being activatory at low concentration, and inhibitory at high concentration. The activatory effect is proposed to result from PIP_2_ being hydrolyzed to PA by phospholipase D2 bound to the C-terminus of TREK channels with the increase in local concentration of PA directly activating TREK-1 (Comoglio *et al*., 2014); by contrast, the inhibitory effect is proposed to result from PIP_2_ competing PA out of its activatory site implying that PA may be the main channel activator, not PIP_2_ (Cabanos *et al*., 2017).

Importantly, our results are consistent with mutagenesis and other functional studies as well as with biochemical studies of lipid binding. The putative PIP_2_ binding site we identify in the up state includes residues R297, K301 and K304 which have previously been implicated in PIP_2_ binding (Chemin *et al*., 2005*b*). Also, the PIP_2_ binding site we identify in the down state involves K304 and R311, and mutation of R311 also affects channel activity in a way that is consistent with its role in PIP_2_ binding (Chemin *et al*., 2005*b*). Our results therefore support the idea that PIP_2_ directly interacts with this region of the proximal C-terminus (Chemin *et al*., 2005*b*). Furthermore, our simulation proposes that the PIP_2_ binding conformation of the channel may change as it transits from the down to the up state.

We also show that PIP_2_ has a lower binding affinity to the down state than to the up state and our atomistic simulations indicate that in the presence of PIP_2_, the down state begins to transit towards the up state. This model is consistent with other studies (McClenaghan *et al*., 2016; Proks *et al*., 2021) and computational electrophysiology studies also suggest that the down state of TREK-2 is less active than the up state (Brennecke & de Groot, 2018). This is also supported by single channel studies of the related TRAAK channel which suggest that the up state is more conductive (Brohawn *et al*., 2019). Thus, we propose that PIP_2_ activates TREK-1 by promoting a transition from the down state to an up state that is capable of supporting a higher degree of channel activity. This change involves an expansion of the distance between M2 and M4 and is accompanied by changing the coordination of PIP_2_ from R311 to K304 in the PIP_2_ binding site.

We have also assessed the affinity of different anionic phospholipids in the PIP_2_ binding site of TREK-1. Overall, the binding affinity of anionic phospholipids is much lower in the down state than in the up state. Our simulations also shows that PA affinity to the channel is approximately four-fold lower than for PIP_2_ in the up state. whereas this affinity is almost negligible in the down state. This suggests that PA may only bind and activate the channel when it is in the up state with PIP_2_ already bound to the channel though the mechanism involved remains unclear.

Overall, these simulations provide additional mechanistic insight and understanding of the conformational dynamics of TREK-1 and its activation by PIP_2_ as well as by other anionic phospholipids. The results also provide a better understanding of the different mechanisms of state-dependent lipid activation of TREK channels and the complex allosteric processes involved in their polymodal activation.

## Materials and methods

### CG simulation set-up

The TREK-1 amino acid sequence was taken from UniProt id O95069. The sequence was then aligned to TREK-2 structure (PDB entry: 6CQ6 – up state) (Lolicato *et al*., 2017); (PDB entry: 4XDJ – down state) (Dong *et al*., 2015) using SWISS-MODEL to generate two all-atom structures (Waterhouse *et al*., 2018). We generated a two-fold symmetry structure for each conformational state of TREK-1 by transplanting the structure of one subunit onto another using PyMOL (Schrodinger LLC, 2015). As the down state does not have an extended C-terminus structure, we extended this segment of TREK-1 by superimposing the α-helical structure based on the subunit A from 6CQ6. The structure was converted from atomistic to coarse-grained using *martinize*.py (Monticelli *et al*., 2008). In this study, all coarse-grained simulations were calculated using the Martini2.3 forcefield. The protein is then placed in the periodic simulation box at a minimum distance of 15 Å from the box edge in all directions.

Palmitoyl-2-oleoylphosphatidylcholine (POPC) is a common lipid headgroup in a mammalian plasma membrane (Casares *et al*., 2019). We use palmitoyl oleyl (PO) as the lipid tail for all lipid headgroups (PIP_2_, PI4P, PI, PG, PA, PS). Bilayers were assembled around the TREK-1 and flooded with the coarse-grained water particles using *insane*.py, with a similar area per lipid between both the upper and the lower leaflet (Wassenaar *et al*., 2015). The bilayers have the ratio of POPC:POPS:PIP_2_ at 1:0:0 for the upper leaflet and 15:3:2 for the lower leaflet. We added POPS as PS is localized exclusively in the cytoplasmic leaflet and accounts for 13-15% of anionic phospholipids in the human cerebral cortex. The CG model of TREK-1 was aligned perpendicular to the bilayer. 0.15 mM NaCl was then generated using *genion* to replace water particles with sodium (Na^+^) and chloride (Cl^−^) ions. Together, each simulation box has a size of 13×13×13 nm^3^, containing ~19,000 CG beads (Abraham *et al*., 2015).

### CG-MD simulations

Simulations were run with GROMACS 2020.1, using the V-rescale thermostat for temperature coupling (Bussi *et al*., 2007) and the Parrinello-Rahman barostat for semi-isotropic pressure (Parrinello & Rahman, 1981). The temperature was coupled to 310 K, and the pressure is kept at 1 bar in xy direction. We applied position restraints force of 1000 kJ mol^−1^ nm^−2^ to the backbone beads to maintain their crystal structures. Energy minimization was carried out using the steepest descents algorithm with an energy cap of 1000 kJ mol^−1^ nm^−2^. The system was then equilibrated for 100 ps with the Berendsen thermostat algorithm (Berendsen *et al*., 1984). For lipid-binding site analysis, each production MD was performed for 10 μs with 20 fs timesteps. The binding sites were then assessed using *pylipid*.py (Song *et al*., 2021). The interactions were counted when the distance between amino acid residue and lipid headgroup was shorter than 0.6 nm and stopped counting when further than 0.8 nm.

### CG-PMF calculations

Snapshots of the final frame were taken from CG-MD simulations for the PMF calculation. TREK-1 and a single PIP_2_ in the target binding site were isolated using PyMOL (Schrodinger LLC, 2015) and re-assembled into POPC bilayers using insane.py (Wassenaar *et al*., 2015). The system was energy minimized and equilibrated with PIP_2_ and the protein backbone restrained for 15 ns. The system was then equilibrated further without PIP_2_ restrained for an additional 15 ns. All position restraints were carried out at 1000 kJ mol^−1^ nm^−2^. To maintain a structural consistency with the crystal structures the protein was then restrained throughout the PMF calculation. PIP_2_ was then pulled from the binding site using steered MD along the collective variable (CV). We defined the collective variable as distances between the PIP_2_ headgroup and reference points. We defined our reference points as the backbone particle of the residue near the PIP_2_ binding site (D294 for the up state, R297 for the down state). To pull the headgroup, we applied an elastic force (1000 kJ mol^−1^ nm^−2^) to pull the PIP_2_ headgroup along the collective variable as described. During our umbrella sampling, we sampled C.V.s with 0.5 Å spacing for optimal histogram overlap. These snapshots are used to initialize independent MD simulations where umbellar potential with the force of 1000 kJ mol^−1^ nm^−2^ was applied to the PIP_2_ headgroup in each window. MD simulations were performed for 1.2 μs to allow for convergence, and the first 200 ns was removed from each window as equilibration. These windows were then used to generate 1D energy landscapes using the weighted histogram analysis method (WHAM) from the *gmx wham* tool with 200 Bayesian bootstraps (Hub *et al*., 2010).

### CG-FEP

The bound positions of PIP_2_ were taken from the PMF simulations and used as input for the FEP calculations. The PIP_2_ headgroup was alchemically transformed into other lipid headgroups (PI4P, PI, PG, PS, PA) using an approach similar to our previous study (Pipatpolkai *et al*., 2020). This was done in chemical space with λ as a coordinate parameter. All transformations were performed separately in both bound states and in bulk bilayer to create complete thermodynamic cycles.

During transformations, CG beads were altered to change PIP_2_ into other lipids. Some beads were transformed into dummies with no interaction properties or different beads with properties matching the target. These alteration choices were made manually by aligning PIP_2_ with other lipids structures. Each transformation was performed in steps where λ was slowly changed from 0 (PIP_2_) to 1 (other lipids). As in this report, each transformation was split into 15 windows where Coulombic and Van der Waals interactions were transformed separately, with soft-core parameters (α = 0.5 and σ = 0.3) for both interactions, similar to our previous study (Corey *et al*., 2021). Coulombic interactions’ λ parameter was perturbed linearly in the first 9 windows and converge to one at the end (λ = 0.00, 0.10, 0.20, …, 1.00), while Van der Waals interactions’ λ parameter was perturbed linearly started from the sixth window toward the final window (λ = 0.00 … 0.00, 0.10, 0.20, …, 0.90, 1.00). Energy minimizations were performed with the steepest descent algorithm for 200 steps. 15 independent production simulations with randomized initial velocities were then run using leap-frog stochastic dynamics integrator to 12 ns per window, where the first 2 ns was discarded as equilibration. Using the *alchemical-analysis* software package, we constructed free energy pathways from individual simulations window with Multistate Bennett Acceptance Ratio (MBAR) (Fajer *et al*., 2009).

### Atomistic simulations

The CG-MD simulation snapshots were taken to isolate the channel with PIP_2_ in the most prominent binding site. We generated a position of PIP_2_ by transplanting the position of the PIP_2_ in the binding site onto another subunit using PyMOL. We then re-assembled the PIP_2_-bound TREK-1 into the POPC bilayer using *insane*.py. Energy minimization and equilibration were carried out using the same method used in CG-PMF. The original CG structures of up state and down state TREK-1 were inserted into 100% POPC bilayer using *insane*.py for an unbound TREK-1 simulation. The system was energy minimized using the steepest descent algorithm and equilibrate with protein backbone restrained for 30 ns at 1000 kJ mol^−1^ nm^−2^ using V-rescale thermostat for temperature coupling (Bussi *et al*., 2007) and the Berendsen barostat for semi-isotropic pressure coupling (Berendsen *et al*., 1984).

The up state and down state coarse-grained structures were converted to atomistic structures using *cg2at*.py (Vickery & Stansfeld, 2021). We inserted potassium (K^+^) ions from the crystal structures into the selectivity filter and converted sodium (Na^+^) ions into potassium (K^+^) ions. Thus, our system in atomistic simulation is solvated in a 0.15 KCl solution. Another energy minimization was performed to adjust the position of the potassium (K^+^) ions inside the selectivity filter. The system was then energy minimized using steepest-descent algorithm for 5000 steps and then equilibrated for 10 ns with the restraint of 1000 kJ nm^−2^ mol^−1^ on the C_α_ atoms. During the equilibration, all systems (both with and without PIP_2_) were coupled with V-rescale thermostat to maintain simulation temperature at 310 K (Bussi *et al*., 2007) and the Berendsen barostat for semi-isotropic pressure coupling (Berendsen *et al*., 1984). Three 500 ns production simulations for each state were performed with different randomized initial velocities. During the production run, V-rescale thermostat is used to maintain the simulation temperature at 310 K (Bussi *et al*., 2007) and the Parrinello-Rahman barostat for semi-isotropic pressure at 1 bar in xy direction (Parrinello & Rahman, 1981). All atomistic simulations and analyses (Root mean square fluctuation, Hydrogen bonding, principal component analysis and distance calculation) were performed using GROMACS 2020.1 with CHARMM36 forcefields (Huang & Mackerell, 2013; Abraham *et al*., 2015; Huang *et al*., 2017). 95% confidence interval from Root mean square fluctuation (RMSF) analysis from each simulation are calculated using 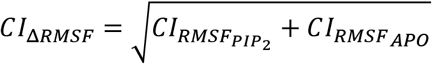. We assessed the amino acid residues that interact with the anionic headgroup of the PIP2 using *pylipid*.py with the cut-off distance of 0.4 nm.

## Acknowledgements

This work was supported by grants from the Biotechnological and Biological Sciences Research Council to S.J.T and by an OXION Programme Studentship from the Wellcome Trust to T.P. A.P. was funded by a Royal Thai Government Scholarship. We also thank members of the Tucker, Sansom, Stansfeld and Biggin groups for their helpful comments during the development of this project.

## Supplementary Figure Legends

**Figure S1.**
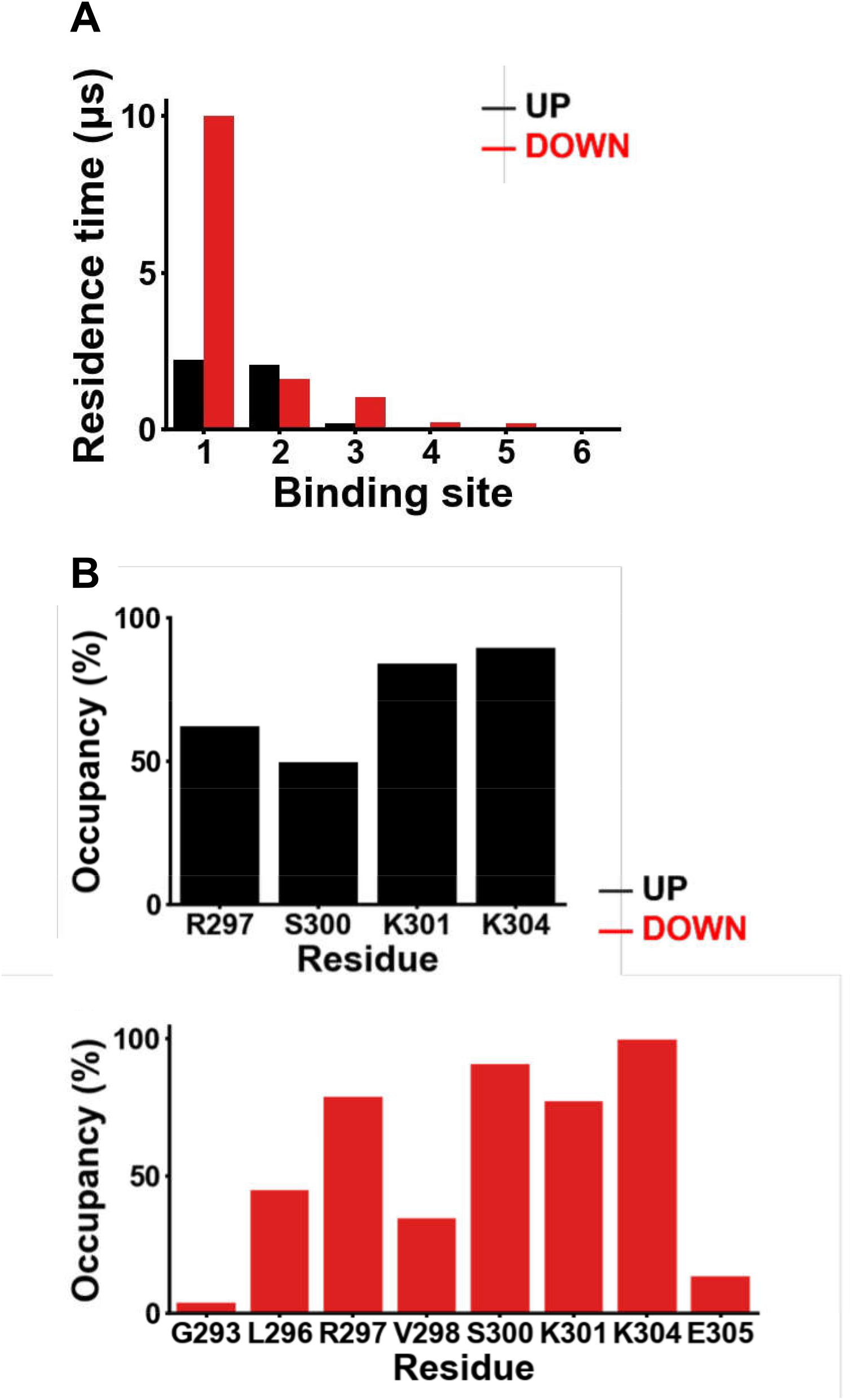
Residence time and occupancy of PIP_2_ binding sites from coarse-grained simulations. (**A**) The residence time of PIP_2_ in binding sites in up state (black) and down state (red), ordered by their residence time in the binding site. (**B**) Bar plot of occupancy of residue in the most prominent binding site in up state (black) and down state (red) from 10 μs CG simulation.

**Figure S2.**
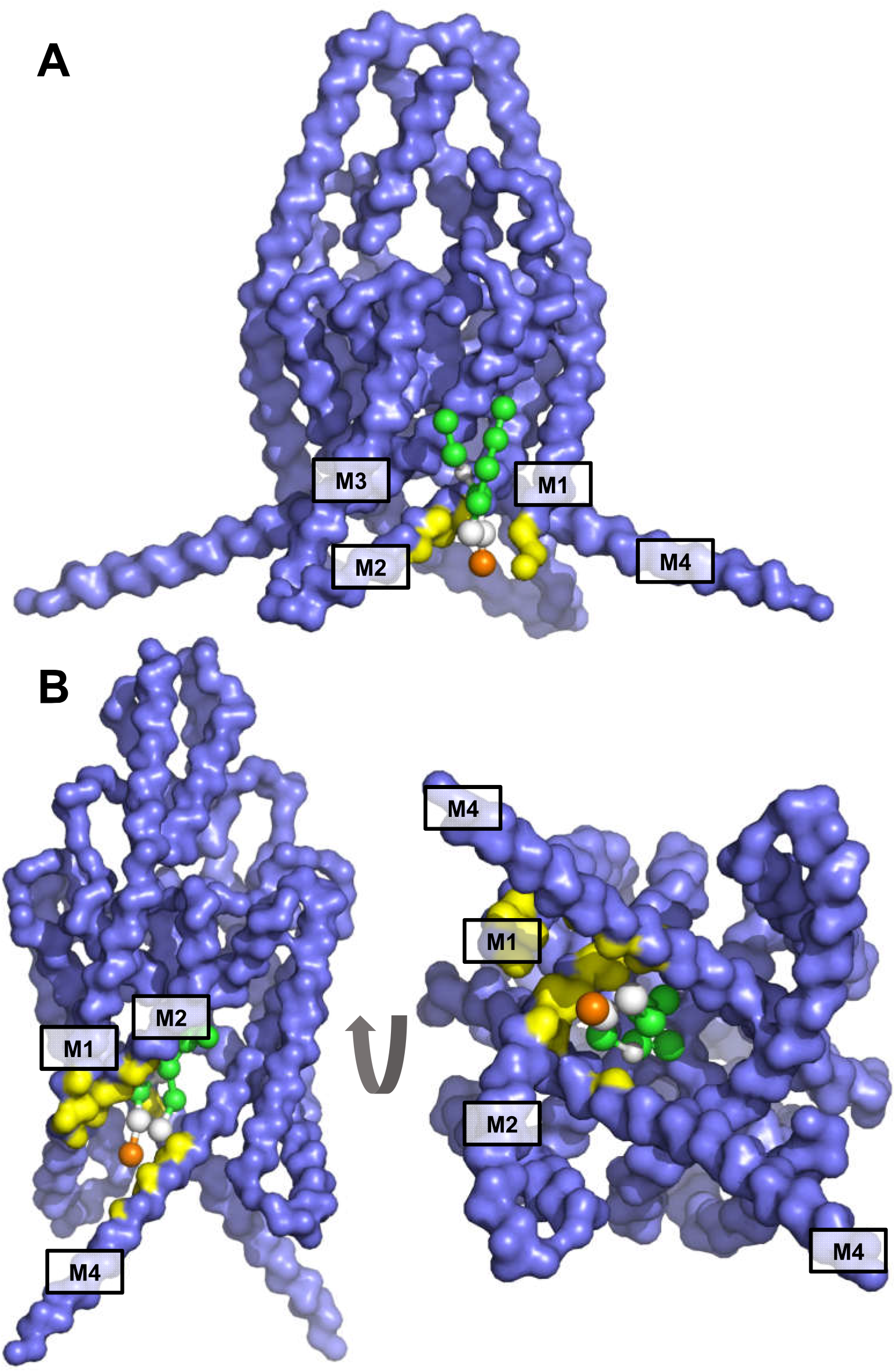
Phosphatidic acid (PA) binding site on the up and down state simulation. The phosphatidic acid binding site with highest occupancy and residence time (yellow) are shown superimposing with (A) up state and (B) down state crystal structures (blue) after a 10 μs CG simulation.

**Figure S3.**
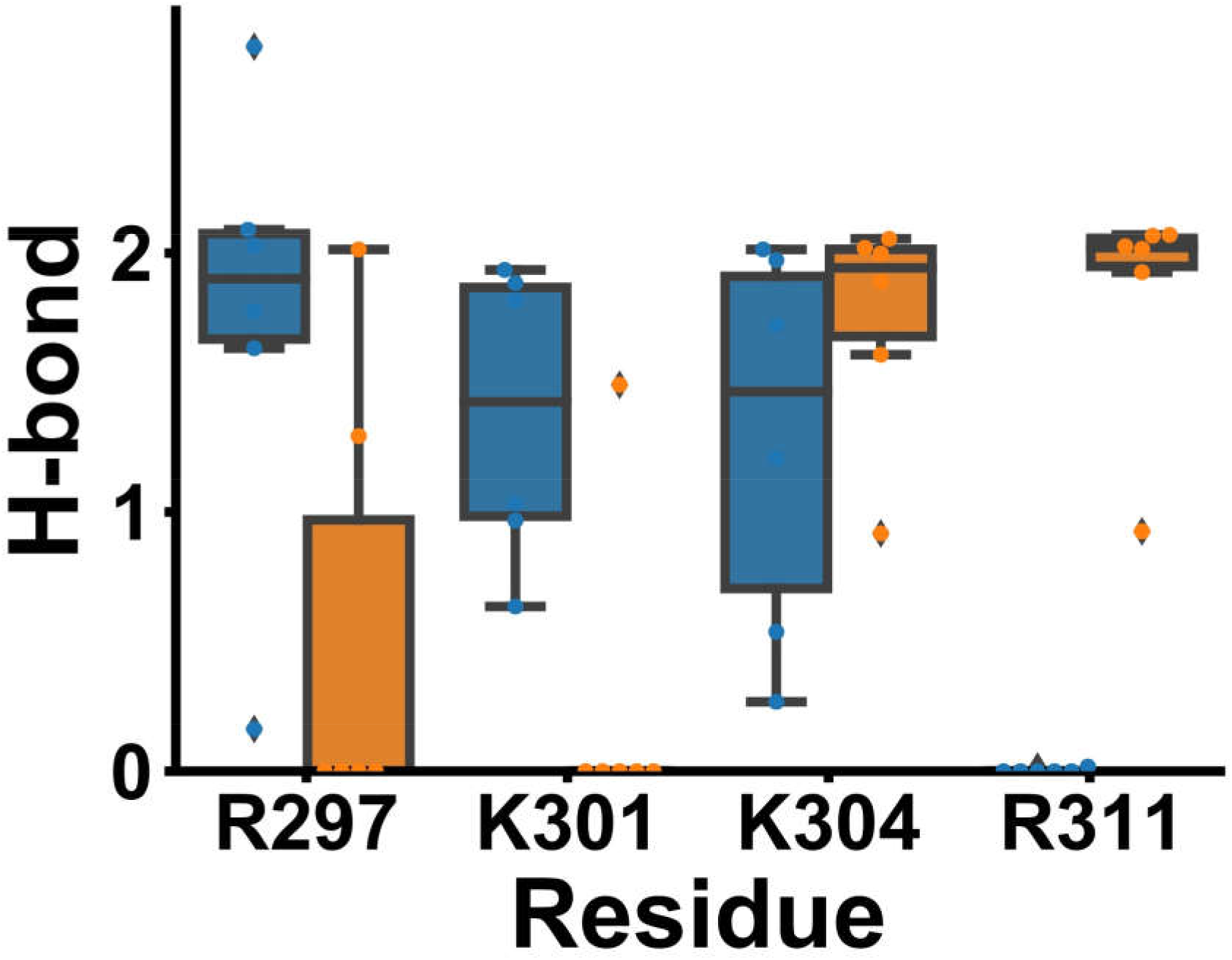
Hydrogen bonding between PIP_2_ and TREK-1 in the up and down states. Median number of hydrogen bonds formed from the last 100 ns of the atomistic simulation of TREK-1 in the up state (blue) or down state (orange). Each plot contains six data points from three simulations and each subunit of the TREK-1 channel. The error bar represents the inter-quartile range of the distribution.

**Figure S4.**
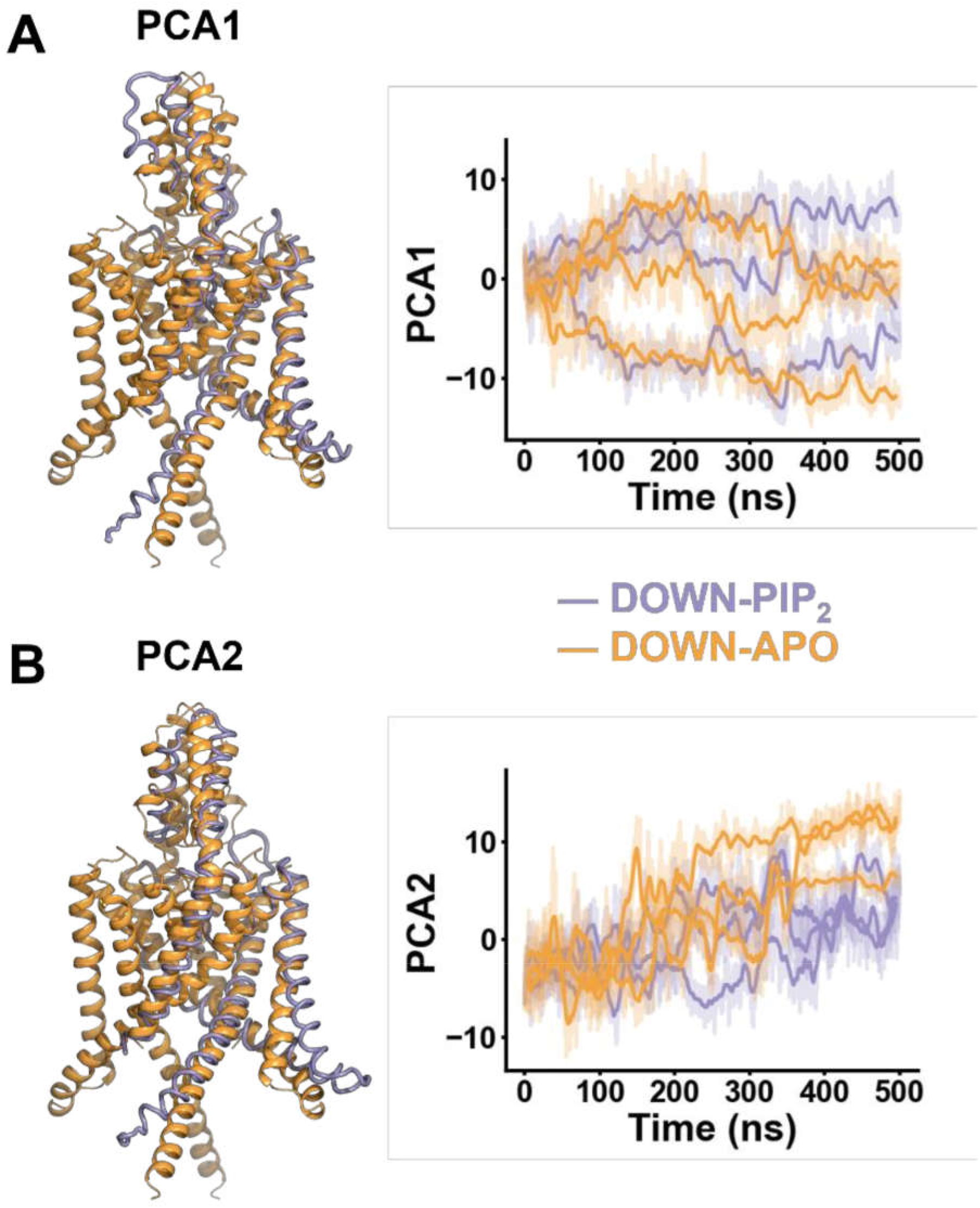
Principle component analysis of PIP2-induced movement in TREK-1. The principal components are calculated from three simulations of TREK-1 channel in the down state containing PIP_2_. (**Left**) The backbone of the protein before the simulation (orange) is aligned to the interpolation of the of the most extreme projection along (**A**) the first principle and (**B**) the second principal component (purple) calculated from the simulation. (**Right**) The magnitude of the conformational change along the vector describing the first and second principal component derived from the simulation over 500 ns (n = 3). The simulations were conducted with PIP_2_ (purple) an without PIP_2_ (orange).

**Fig S5.**
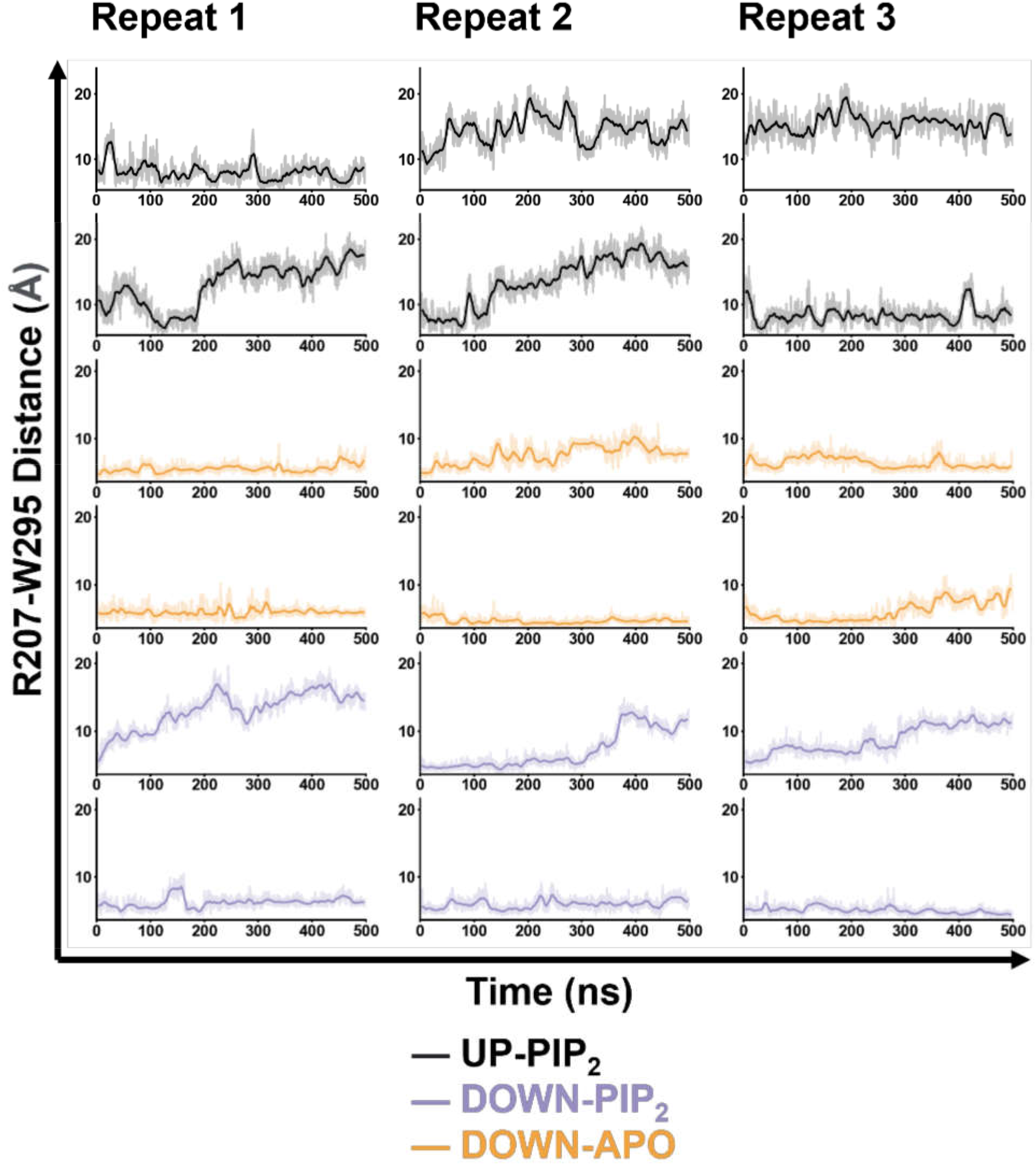
R207-W295 (zipper) distance in all simulations. Distances between R207-W295 were calculated in three different simulation set-up (up state with PIP_2_ - black, Down state with PIP_2_ – purple, Down state without PIP_2_ - orange). Each measurement represents the distance within each subunit of the TREK-1 channel.

